# Therapeutic inhibition of keratinocyte TRPV3 sensory channel by local anesthetic dyclonine

**DOI:** 10.1101/2020.09.21.306316

**Authors:** Qiang Liu, Juan Hu, Jin Wang, Xin Wei, Conghui Ping, Yue Gao, Chang Xie, Ye Yu, Dongdong Li, Jing Yao

**Affiliations:** State Key Laboratory of Virology, Hubei Key Laboratory of Cell Homeostasis, College of Life Sciences, Frontier Science Center for Immunology and Metabolism, Wuhan University, Wuhan, Hubei 430072, China; School of Basic Medicine and Clinical Pharmacy, China Pharmaceutical University, Nanjing, 211198, China; Sorbonne Université, Institute of Biology Paris Seine, Neuroscience Paris Seine, CNRS UMR8246, INSERM U1130, UPMC UM119, Paris 75005, France

**Author notes:** Address correspondence to: Dr. Jing Yao, State Key Laboratory of Virology, Hubei Key Laboratory of Cell Homeostasis, College of Life Sciences, Frontier Science Center for Immunology and Metabolism, Wuhan University, Wuhan, Hubei 430072, China.

**Keywords:** TRPV3, Dyclonine, Cell death, Pruritus, Skin inflammation

## Abstract

The multimodal sensory channel transient receptor potential vanilloid-3 (TRPV3) is expressed in epidermal keratinocytes and implicated in chronic pruritus, allergy, and inflammation-related skin disorders. Gain-of-function mutations of TRPV3 cause hair growth disorders in mice and Olmsted Syndrome in human. Nevertheless, whether and how TRPV3 could be therapeutically targeted remains to be elucidated. We here report that mouse and human TRPV3 channel is selectively targeted by the clinical medication dyclonine that exerts a potent inhibitory effect. Accordingly, dyclonine rescued cell death caused by gain-of-function TRPV3 mutations and suppressed pruritus symptoms in vivo in mouse model. At the single-channel level, dyclonine inhibited TRPV3 open probability but not the unitary conductance. By molecular docking and mutagenesis, we further uncovered single residues in TRPV3 pore region that could toggle the inhibitory efficiency of dyclonine. The functional and mechanistic insights obtained on dyclonine-TRPV3 interaction will help to conceive updated therapeutics for skin inflammation.

## Introduction

Transient receptor potential (TRP) channels belong to a family of calcium-permeable and nonselective cation channels, essential for body sensory processing and local inflammatory development (Clapham, 2003). As a polymodal cellular sensor, TRPV3 channel is abundantly expressed in skin keratinocytes (Peier et al., 2002; Xu et al., 2002; Chung et al., 2004a) and in cells surrounding the hair follicles (Cheng et al., 2010). TRPV3 integrates a wide spectrum of physical and chemical stimuli (Luo and Hu, 2014). TRPV3 is sensitive to innocuous temperatures above 30-33 °C and exhibits an increased response at noxious temperature (Xu et al., 2002; Chung et al., 2005). Natural plant products such as camphor (Moqrich et al., 2005), carvacrol, eugenol, thymol (Xu et al., 2006), and the pharmacological compound 2-aminoethoxydiphenyl borate (2-APB) (Chung et al., 2004b; Colton and Zhu, 2007) also activate TRPV3. In addition, TRPV3 is directly activated by acidic pH from cytoplasmic side (Gao et al., 2016).

Mounting evidence implicates TRPV3 channel in cutaneous sensation including thermal sensation (Chung et al., 2004a), nociception (Huang et al., 2008), and itch (Yamamoto-Kasai et al., 2012). They also participate in the maintenance of skin barrier, hair growth (Cheng et al., 2010) and wound healing (Yamada et al., 2010; Aijima et al., 2015). Recently, the dysfunction of TRPV3 channels has come to the fore as a key regulator of physio- and pathological responses of skin (Ho and Lee, 2015). In rodents, the Gly573Ser substitution in TRPV3 renders the channel spontaneously active and caused a hairless phenotype in DS-Nh mice and WBN/Kob-Ht rats (Asakawa et al., 2006). DS-Nh mice also develop severe scratching behavior and pruritic dermatitis. TRPV3 dysfunction caused by genetic gain-of-function mutations or pharmaceutical activation has been linked to human skin diseases including genodermatosis known as Olmsted syndrome (Lin et al., 2012; Agarwala et al., 2016) and erythromelalgia (Duchatelet et al., 2014). Furthermore, TRPV3-deficient mice give rise to phenotypes of curly whiskers and wavy hair coat (Cheng et al., 2010). Conversely, hyperactive TRPV3 channels expressed in human outer root sheath keratinocytes inhibit hair growth (Borbiro et al., 2011). While being implicated in a variety of skin disorders, whether and how TRPV3 could be therapeutically targeted remains to be elucidated. It is thus desirable to identify and understand clinical medications that can selectively target TRPV3 channels.

As a clinical anesthetic, dyclonine is characterized by rapid onset of effect, lack of systemic toxicity, and a low index of sensitization (Florestano and Bahler, 1956). Its topical application rapidly relieves itching and pain in patients, by ameliorating inflamed, excoriated and broken lesions on mucous membranes and skin (Morginson et al., 1956). Accordingly, dyclonine is used to anesthetize mucous membranes prior to endoscopy (Formaker et al., 1998). The clinical scenario targeted by dyclonine treatment echoes the pathological aspects of TRPV3-related skin disorders, suggesting that the therapeutic effects of dyclonine might involve its interaction with TRPV3 sensory channel.

Here, using a multidisciplinary approach combining electrophysiology, genetic engineering and ultrafast local temperature control, we show that mouse and human TRPV3 channel was selectively targeted by dyclonine. It dose-dependently inhibited TRPV3 currents in a voltage-independent manner and rescued cell death caused by TRPV3 gain-of-function mutation. In vivo, dyclonine indeed suppressed the itching/scratching behaviors induced by TRPV3 channel agonist carvacrol or pruritogen histamine. At single-channel level, dyclonine reduced TRPV3 channel open probability without altering the unitary conductance. We also identified molecular residues that were capable of either eliminating or enhancing the inhibitory effect of dyclonine. These data demonstrate the selective inhibition of TRPV3 channel by dyclonine, supplementing a molecular mechanism for its clinical effects and raising its potential to ameliorate TRPV3-associated disorders.

## Results

### Inhibition of TRPV3 currents by dyclonine

We first examined the effect of dyclonine on TRPV3 activity induced by the TRPV channel agonist 2-APB (100 μM). Whole-cell currents were recorded at a holding potential of −60 mV in HEK 293T cells expressing mouse TRPV3. Because TRPV3 channels exhibit sensitizing properties upon repeated stimulation (Chung et al., 2004c), we examined the effect of dyclonine after the response had stabilized following repetitive application of 2-APB (Fig. 1A). The presence of 5 μM and 10 μM dyclonine significantly inhibited TRPV3 currents response to 30 ± 2% and 15 ± 3% of control level, respectively. After washing out of dyclonine, 2-APB evoked a similar response to the control level, indicating the blocking effect of dyclonine is reversible (Fig. 1A-B). We repeated the experiments with different doses of dyclonine. The dose-response curve indicates that dyclonine inhibited TRPV3 currents in a concentration-dependent manner with an IC_50_ of 3.2 ± 0.24 μM (*n* = 6, Fig. 1C).

**Figure 1.**
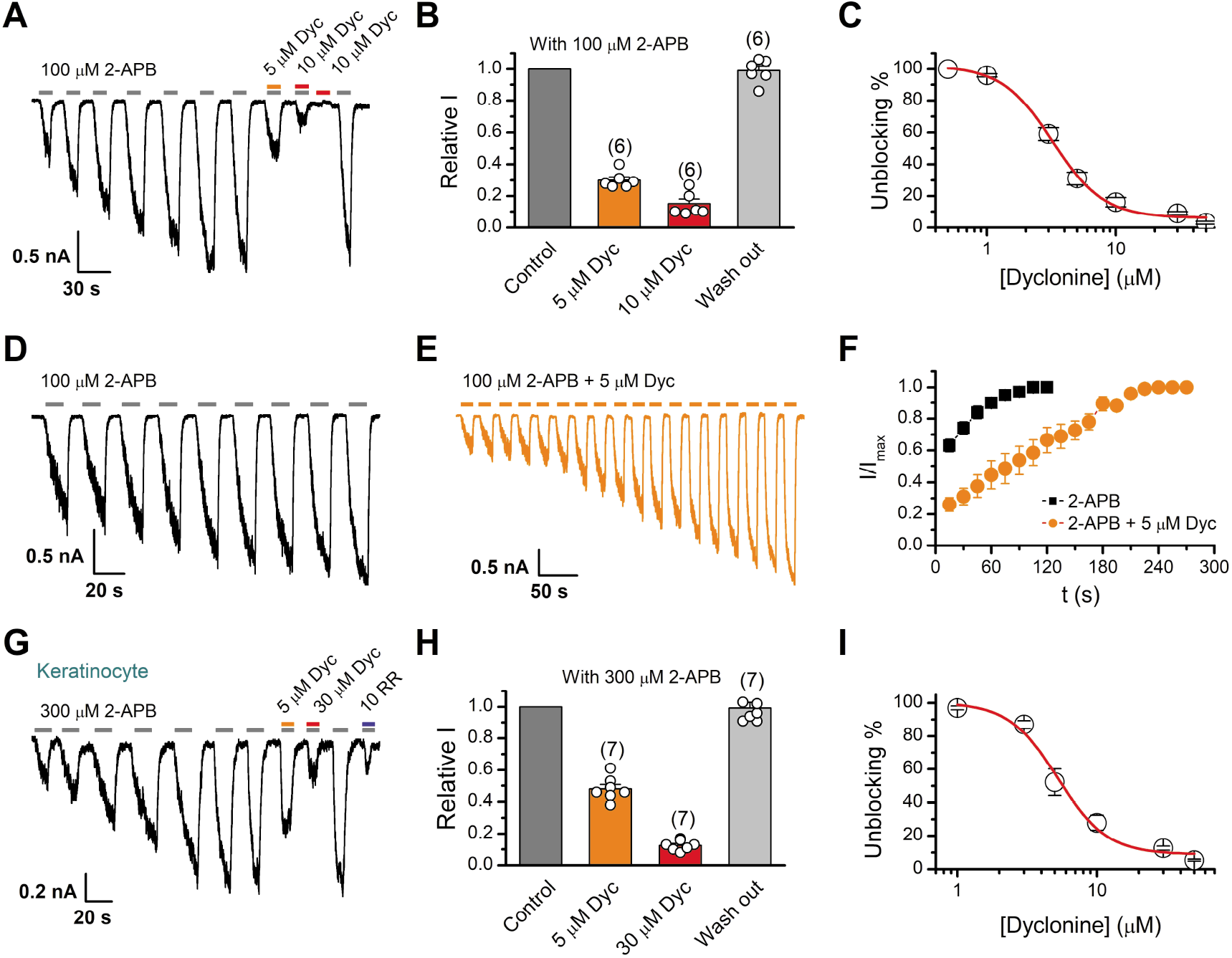
Inhibition of TRPV3 currents by dyclonine. (**A**) Inhibition of 2-APB-evoked currents by dyclonine (Dyc) in a representative HEK 293T cell expressing mouse TRPV3. After sensitization by repeated application of 100 μM 2-APB, the cell was exposed to 5 or 10 μM dyclonine with 100 μM 2-APB or 10 μM dyclonine only as indicated by the bars. Membrane currents were recorded in whole-cell configuration, and the holding potential was −60 mV. (B) Summary of relative currents elicited by 100 μM 2-APB in the presence of 0, 5 or 10 μM dyclonine. Numbers of cells are indicated in parentheses. (**C**) The dose-response curve for dyclonine inhibition of TRPV3 currents was fitted by Hill’s equation (IC_50_ = 3.2 ± 0.24 μM and n_H_ = 2.2 ± 0.32, *n* = 6). (**D-E**) Representative whole-cell recordings for the sensitization of TRPV3 currents elicited by repeated applications of 100 μM 2-APB in the absence (**D**) and presence (**E**) of 5 μM dyclonine. (**F**) Time courses toward the peak currents elicited by repeated application of 100 μM 2-APB with or without dyclonine *(n* = 9). Currents were normalized to the maximum values after sensitization. (**G**) The 2-APB-evoked inward currents were reversibly inhibited by dyclonine in primary mouse epidermal keratinocytes. Representative inward currents were firstly activated by repeated application of 300 μM 2-APB at the holding potential of −60 mV, and then inhibited by 5 or 30 μM dyclonine or 10 μM Ruthenium Red (RR) as indicated. (**H**) Summary of relative currents elicited by 300 μM 2-APB with or without dyclonine. (**I**) Dose dependence of dyclonine effects on TRPV3 currents in cultured keratinocytes. The solid line corresponds to a fit by Hill’s equation with IC_50_ = 5.2 ± 0.71 μM and n_H_ = 2.4 ± 0.75 (*n* = 6).

TRPV3 channel in physiological conditions has a low level of response to external stimuli, which is augmented during the sensitization process (i.e., repetitive stimulations, Fig. 1A). In contrast, excessive up-regulation of TRPV3 activity impairs hair growth and increases the incidence of dermatitis and pruritus in both humans and rodents. To determine whether dyclonine affects the process of TRPV3 sensitization, TRPV3-expressing cells were repeatedly exposed to 100 μM 2-APB without or with 5 μM dyclonine (Fig. 1D, E). TRPV3 currents evoked by 2-APB alone took ~8 repetitions to reach full sensitization level (Fig. 1D). The presence of dyclonine significantly slowed down this process, requiring ~16 repetitions to reach the current level of full sensitization (Fig. 1E-F). As expected, dyclonine also reduced the initial TRPV3 current (31.12 ± 2.86 pA/pF, v.s. 86.43 ± 5.9 pA/pF without dyclonine; p < 0.001; *n* = 9 per condition).

As TRPV3 is highly expressed in keratinocytes, we further determined the inhibitory effect of dyclonine in primary mouse epidermal keratinocytes. After stabilizing the channel current by repeated application of 2-APB, we tested the inhibitory effect of 5 μM and 30 μM dyclonine (Fig. 1G). On average, TRPV3 currents were reduced to 52 ± 7% and 13 ± 0.01% of control level by 5 μM and 30 μM dyclonine, respectively (Fig. 1H), reaching the similar level of inhibition by the wide-spectrum TRP channel blocker ruthenium red (RR, Fig. 1G). From the dose-response curve (Fig. 1I), the IC_50_ of dyclonine was assessed to be 5.2 ± 0.71 μM, with a Hill coefficient of n_H_ = 2.4 ± 0.75 (*n* = 7). Thus, dyclonine effectively suppresses the activity of endogenous TRPV3 channels in mouse keratinocytes.

### Dyclonine is a selective inhibitor of TRPV3 channel

Next, we compared the inhibitory effect on TRPV3 of dyclonine to its impact on other TRP channels. Mouse TRPV1, TRPV2 and TRPM8 channels were expressed in HEK 293T cells and respectively activated by capsaicin, 2-APB and menthol. We observed that 10 μM dyclonine exhibited little inhibition on TRPV1, TRPV2 and TRPM8, but potently inhibited TRPV3 channel (Fig. 2A). The corresponding reduction in current amplitude was 2 ± 1% for TRPV1, 6 ± 1% for TRPV2, 3 ± 2% for TRPM8, compared with 84 ± 1% inhibition of TRPV3 current (Fig. 2B). By applying a series of dyclonine concentrations, we derived dose-response curves (Fig. 2C). The corresponding IC_50_ values of dyclonine for inhibiting TRPV1, TRPV2 and TRPM8 channels (337.4 ± 19.4 μM, 31.1 ± 2.7 μM and 81.8 ± 12.7 μM, respectively) were one or two orders of magnitude higher than that for TRPV3 inhibition (3.2 ± 0.24 μM), indicating that dyclonine represents a selective inhibitor of TRPV3 channel.

**Figure 2.**
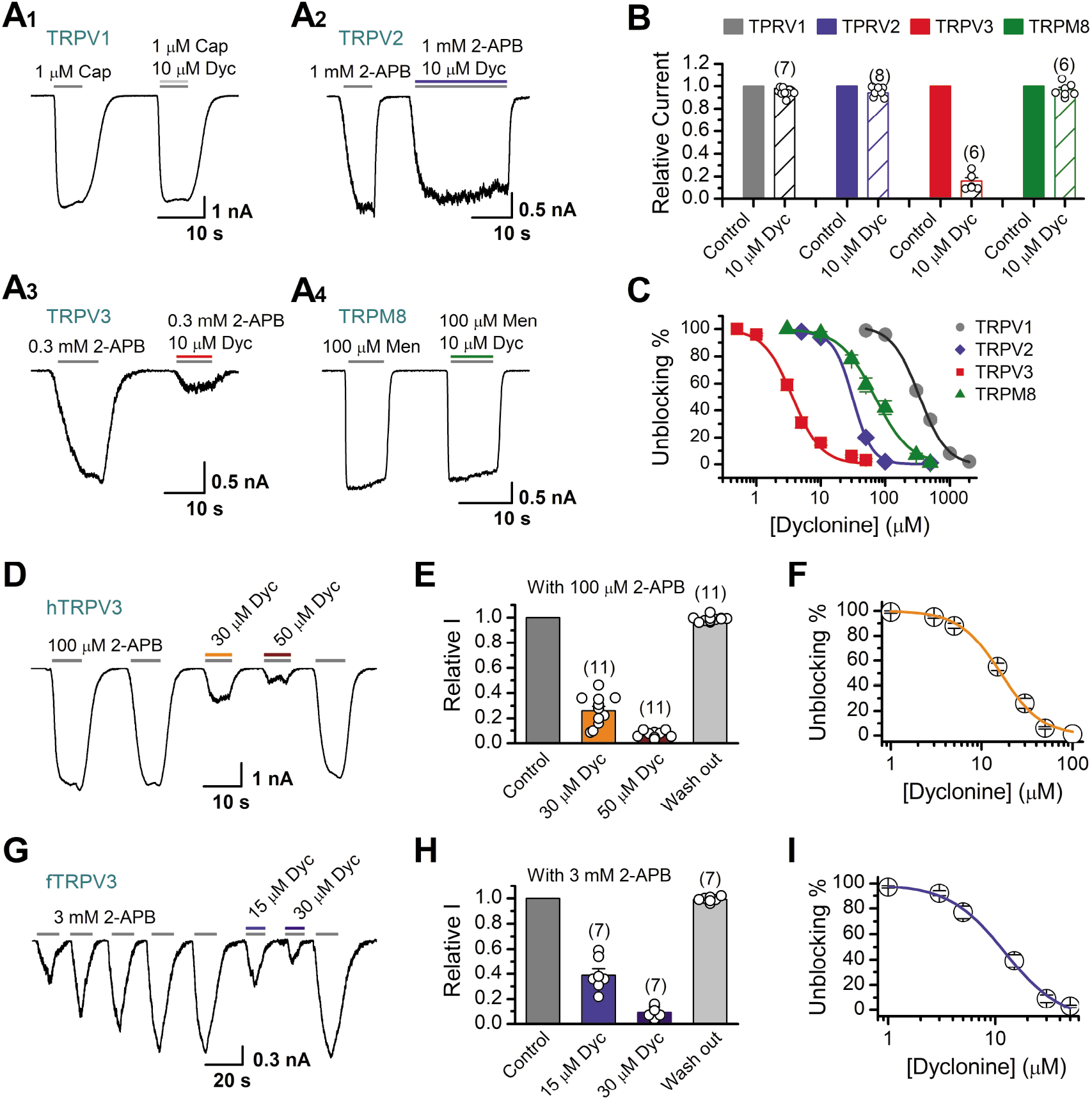
Selective inhibitory effect of dyclonine on TRPV3. (**A**) Representative inward current traces from whole-cell voltage-clamp recordings show the inhibitory effects of 10 μM dyclonine on TRPV1 (**A_1_**), TRPV2 (**A_2_**), TRPV3 (**A_3_**), or TRPM8 (**A_4_**) channels (Cap, capsaicin; Men, menthol). Bars represent duration of drug application. (**B**) Summary of relative currents before and after dyclonine (10 μM) treatment. Numbers of cells are indicated in parentheses. (**C**) Dose-response curves of dyclonine for inhibition of indicated ion channel currents. Solid lines represent fits by Hill equation, with IC_50_ = 337.4 ± 19.4 μM and n_H_ = 2.0 ± 0.31 for TPRV1 (*n* = 7), IC_50_ = 31.1 ± 2.7 μM and n_H_ = 2.9 ± 0.50 for TPRV2 (*n* = 8), IC_50_ = 81.8 ± 12.7 μM and n_H_ = 1.2 ± 0.20 for TRPM8 (*n* = 6). For comparison, the dose-response curve of TRPV3 channel from Fig. 1C is displayed in red with IC_50_ = 3.2 ± 0.24 μM and n_H_ = 2.2 ± 0.32 (*n* = 6). (**D**) Suppression of 2-APB-evoked currents by dyclonine in a hTRPV3-expressing HEK293T cell. Representative inward current trace shows the reversible block effect of dyclonine (30 and 50 μM) at the holding potential of −60 mV (**E**) Summary of inhibition of hTRPV3 by dyclonine. Membrane currents were normalized to the responses elicited by the saturated concentration of 2-APB (100 μM) alone. (**F**) Dose-response curve for dyclonine on blocking of hTRPV3. Solid line represents a fit to a Hill equation, yielded IC_50_ = 16.2 ± 0.72 μM and n_H_ = 1.91 ± 0.14 (*n* = 11). (**G**) Inhibition of fTRPV3 currents by dyclonine. Representative whole-cell currents at −60 mV in a fTRPV3-expressing HEK293T cell. After sensitization by repeated application of 3 mM 2-APB, the cell was exposed sequentially to 15 and 30 μM dyclonine with 3 mM 2-APB. (**H**) Summary of inhibition of relative currents elicited by 3 mM 2-APB, 3 mM 2-APB with dyclonine 15 or 30 μM. (**I**) Concentration-response curve of dyclonine on the inhibition of fTRPV3 currents. Solid line represents a fit by a Hill equation, with IC_50_ = 12.31 ± 1.6 μM and n_H_ = 1.6 ± 0.34 (*n* = 7). Dyc, dyclonine; hTRPV3, human TRPV3; fTRPV3, frog TRPV 3.

Above results were obtained for mouse TRPV3. We further asked whether the inhibitory effect of dyclonine on TRPV3 is consistent across different species. Similarly, we performed whole-cell recordings in HEK 293T cells expressing human TRPV3 and frog TRPV3, respectively. They were activated to a stable level by repetitive 2-APB stimulation. Addition of dyclonine, indeed, efficiently suppressed the activation of both types of TRPV3 channel (Fig. 2D-I). Dose-response curves for dyclonine inhibition yielded an IC_50_ value of 16.2 ± 0.72 μM for hTRPV3 and 12.3 ± 1.6 μM for fTRPV3, respectively. Therefore, the specific inhibition of TRPV3 by dyclonine is conserved across species.

### Inhibition of TRPV3 by dyclonine is voltage-independent

To obtain a complete description of the inhibitory effect of dyclonine, we next investigated its voltage dependence using a stepwise protocol (Fig. 3A). We measured membrane currents in TRPV3-expressing HEK 293T cells using a Cs^+^-based pipette solution that blocks most outward K^+^ channel current but permits measurement of outward conductance mediated by the nonselective TRPV3 channel. A low-concentration 2-APB (40 μM) activated small voltage-dependent currents with steady-state outward rectification, characteristic of TRPV3 currents in heterologous expression systems (Fig. 3A). Addition of dyclonine in the extracellular solution significantly diminished TRPV3-mediated outward and inward currents (Fig. 3A). By contrast, 10 μM ruthenium red, a broad TRP channel blocker, only inhibited TRPV3-mediated inward currents but enhanced outward currents (Fig. 3A), which is consistent with early report (Cheng et al., 2010). Dyclonine inhibition of both inward and outward currents was further confirmed by the I-V curves derived from pooled data (Fig. 3B). Such inhibition occurred independently of the membrane potential. Together, relative to the wide-spectrum blocker ruthenium red, dyclonine more effectively inhibits TRPV3 channel in a voltage-independent manner.

**Figure 3.**
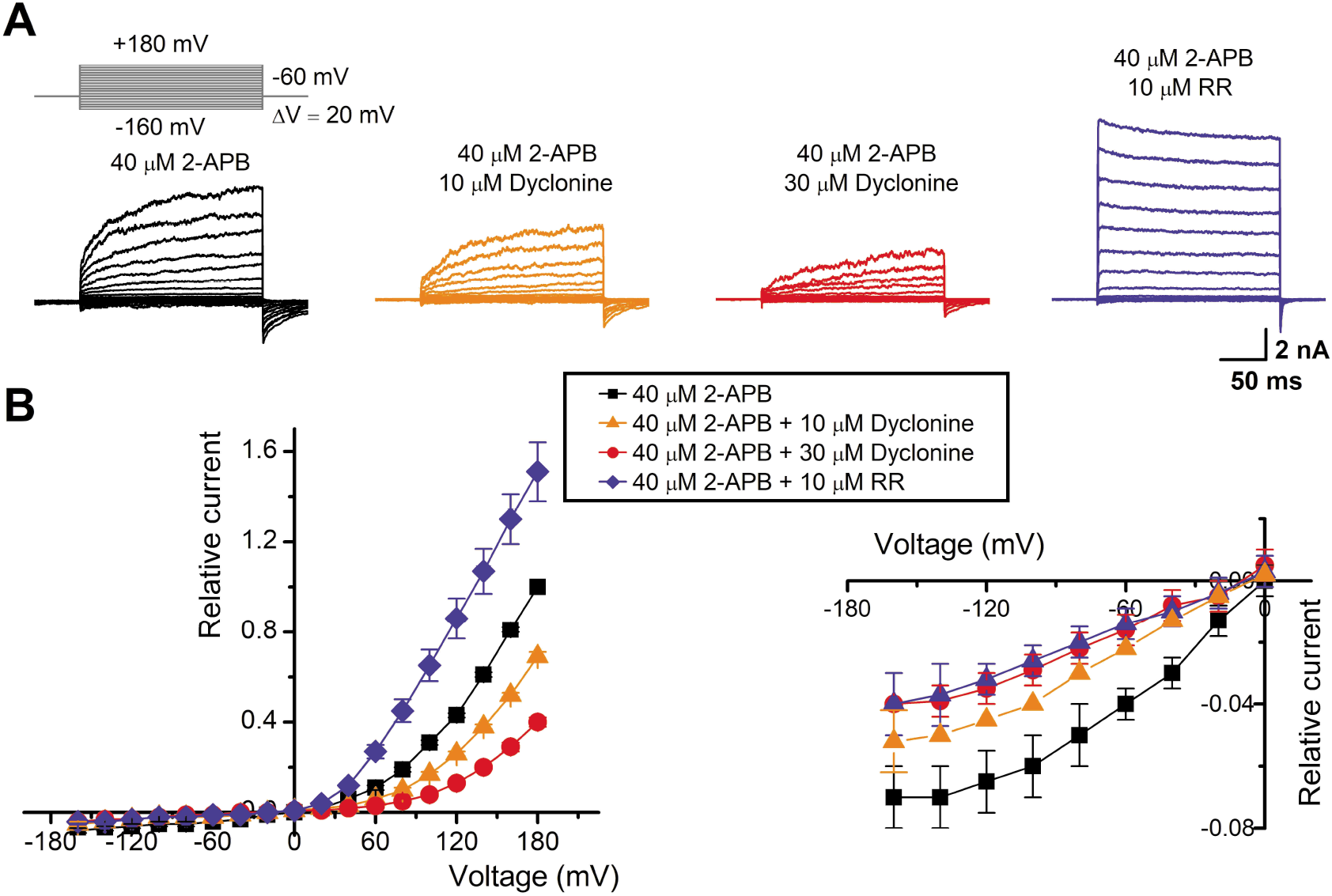
The inhibitory effect of dyclonine on TRPV3 channel is voltage-independent. (**A**) Representative whole-cell currents evoked by voltage steps (*inset*) together with 40 μM 2-APB in the absence and presence of 10, 30 μM dyclonine or 10 μM RR in HEK 293T cells expressing mouse TRPV3. Currents were elicited with 200-ms test pulses ranging from −160 mV to +180 mV in 20-mV increments within the same cells, and the holding potential was −60 mV. Calcium-free standard bath solution and a CsCl-filled recording electrode were used. (**B**) Current-voltage relations for data in (A). Current amplitudes were normalized to the maximum responses at +180 mV in the presence of 40 μM 2-APB. Each point represents mean values (± SEM) from nine independent cells. (*Inset*) The inhibition effects of dyclonine and RR on TRPV3 currents at negative holding potentials is magnified and displayed on the right. Note that dyclonine had an inhibitory effect on TRPV3 currents at both positive and negative potentials, but RR only inhibited TRPV3 channel currents at negative potentials while enhanced TRPV3 currents at positive potentials (left panel, blue trace).

### Inhibition of heat-activated TRPV3 currents by dyclonine

TRPV3 is a thermal sensitive ion channel and has an activation threshold around 30 to 33 °C (Xu et al., 2002). We therefore explored whether the heat-evoked TRPV3 currents can be also inhibited by dyclonine. We employed an ultrafast infrared laser system to control the local temperature near single cells; each temperature jump had a rise time of 1.5 ms and lasted for 100 ms (Fig. 4A). TRPV3, expressed in HEK 293T cells, steadily responded to temperature jumps ranging from 30 to 51 °C (Fig. 4B). As in 2-APB experiments, TRPV3 currents were pre-sensitized to stable level by repeated temperature jumps from room temperature to 52 °C. Application of dyclonine appreciably inhibited TRPV3 thermal currents (Fig. 4B-C). The inhibitory effect of dyclonine was fully reversible, as after its washing out the TRPV3 response recovered to the same level as control condition (Fig. 4C). To determine the concentration dependence of dyclonine inhibition, TRPV3 currents were evoked by a same temperature jump from room temperature to ~52 °C in the presence of 1, 3, 5, 10, 30, and 50 μM dyclonine (Fig. 4D). The IC_50_ of dyclonine on TRPV3 inhibition was assessed to be 14.02 ± 2.5 μM with a Hill coefficient of n_H_ = 1.9 ± 0.54, according to the dose-response curve fitting (Fig. 4E). These results thus indicate that dyclonine dose-dependently suppresses heat-evoked TRPV3 currents.

**Figure 4.**
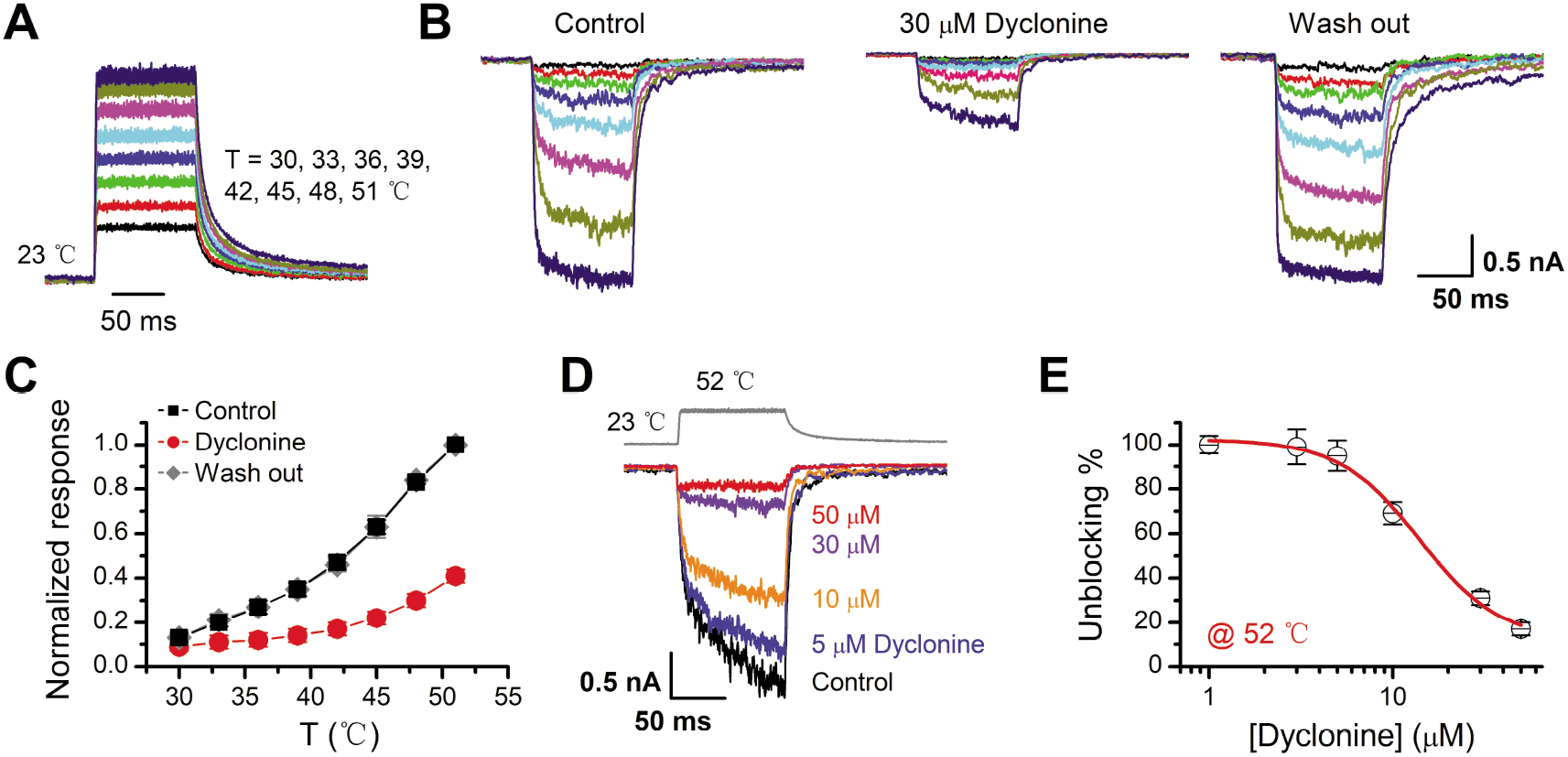
Inhibition of heat-activated TRPV3 currents by dyclonine. (**A**) Temperature pulses generated by infrared laser diode irradiation ranging from 30 to 51 °C, and each pulse had a duration of 100 ms and a rise time of 1.5 ms. (**B**) Effects of dyclonine on heat-activated TRPV3 currents. Heat-evoked current traces were recorded in whole-cell configuration, which were stabilized by sensitization of repeated fast temperature jumps from room temperature to 52 °C. Bath solution with 0 or 30 μM dyclonine was applied by brief perfusion to the patch just before temperature stimulation on the same cells. (**C**) The average plot compares the temperature responses in the absence and presence of 30 μM dyclonine (*n* = 6). Currents were normalized by their maximum responses under control condition, respectively. Note that data from control and washout are superimposed. (**D**) Representative inward currents evoked by a series of identical temperature jumps inhibited by dyclonine in a concentration-dependent manner. The temperature pulse (52 °C) is shown in gray. Holding potential was −60 mV. (**E**) Dose dependence of dyclonine effects on heat-activated TRPV3 currents. The solid line represents a fit to Hill’s equation with IC_50_ = 14.1 ± 2.5 μM and n_H_ = 1.9 ± 0.54 (*n* = 10). All whole-cell recordings were got from TRPV3-expressing HEK 293T cells held at −60 mV.

### Dyclonine inhibited hyperactive TRPV3 mutants and rescued cell death

It has previously been reported that gain-of-function mutations, G573S and G573C, of TRPV3 are constitutively active and their expression causes cell death (Xiao et al., 2008). As TRPV3 channel is effectively inhibited by dyclonine, we explored whether it can rescue the cell death caused by those mutants. We transfected the inducible cDNA constructs encoding respectively the GFP-tagged wild-type TRPV3, G573S, or G573 mutant into T-Rex 293 cells and then applied 20 ng/ml doxycycline to induce the gene expression. Cells were exposed to different pharmacological drugs (dyclonine, 2-APB, 2-APB and dyclonine, or ruthenium red). Cell death was recognized by the narrow and contracted footprints in bright-field images, and the protein expression level meanwhile monitored by GFP fluorescence. As shown in Fig. 5A, massive cell death was seen in cells that expressed G573C and G573S TRPV3 mutants but not those expressing the wild type TRPV3. Addition of dyclonine largely prevented the cell death while not causing change in the expression level of TRPV3 channels (Fig. 5A), indicating that dyclonine decreased the cytotoxicity caused by the gain-of-function mutants. We further performed flow cytometry analysis and observed that the cell death ratio was maintained at low level (4.96 ± 0.87%, *n* = 7) in cells expressing wild-type TRPV3 (Fig. 5B). By contrast, the expression of G573S or G573C mutant significantly increased the cell death ratio to 45.36 ± 5.79% (*n* = 7) and 52.74 ± 4.94% (*n* = 7), that were effectively reduced by dyclonine (50 μM) to 12.45 ± 2.54% (*n* = 7) and 14.98 ± 4.40% (*n* = 7), respectively. The cell-protective effect of dyclonine was mirrored by the general TRP channel blocker ruthenium red (Fig. 5A-C). As expected, activation of TRPV3 channels with the agonist 2-APB caused significant cell death even in cells expressing wild-type channel and exacerbated the cell death in those expressing the mutant channel G573S (Fig. 5C). Application of dyclonine also reversed the cell death caused by 2-APB activation (9.12 ± 1.42% v.s. 43.73 ± 3.46% for wild-type condition, 17.68% ± 5.66% v.s. 53.60 ± 5.88% for G573S, and 13.85% ± 2.49% v.s. 47.91 ± 5.54% for G573C after and before addition of dyclonine). Collectively, these results indicate that dyclonine rescues cell death by selectively inhibiting the excessive activity of TRPV3 channel.

**Figure 5.**
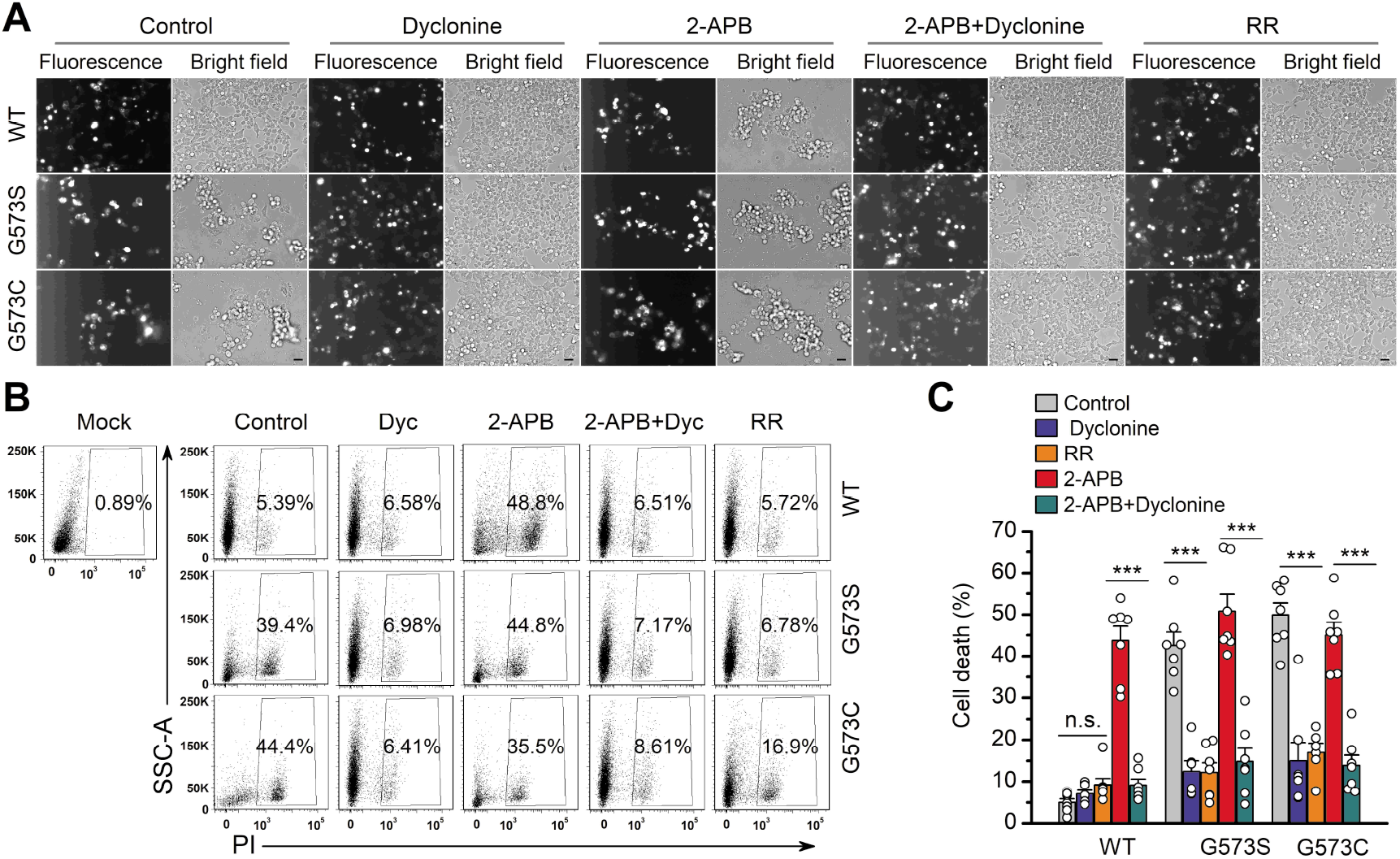
Dyclonine rescued cell death caused by expression of overactive TRPV3 mutant. Bright field and fluorescence images showing the cell survival. The GFP-tagged TRPV3 (wild-type, WT) and two mutants (G573C and G573S) in pcDNA4/TO vector were respectively transfected into T-Rex 293 cells, and then treated with doxycycline (20 ng/ml) for 16 h post-transfection to induce gene expression in the presence of drugs as indicated. Images of cells were taken at 12 h after induction. Scale bar, 50 μm. (**B**) Flow cytometry analysis of the percentage of dead cells. Cells transfected with the desired plasmids are as indicated. After the gene expression induced by doxycycline, the cells were treated with dyclonine (Dyc, 50 μM), 2-APB (30 μM), ruthenium red (RR, 10 μM) or the combination of 2-APB and dyclonine, and then stained with propidium iodide, followed by flow cytometry to analyze cell survival. (**C**) Summary plots of cell death rates under different treatments. Data were averaged from seven independent experiments. *** *P* < 0.0001.

### Dyclonine suppresses acute itch by inhibiting TRPV3 hyperactivity

TRPV3 is highly expressed in skin keratinocytes, whose hyperactivity causes pruritic dermatitis and scratching behavior. We next examined in vivo the therapeutic effect of dyclonine on TRPV3 hyperactivity-caused scratching behavior in mouse model. Itching-scratching behavior was induced by pharmacological activation of TRPV3 channel by a natural compound carvacrol derived from oregano (Cui et al., 2018). The number of scratching bouts was quantified every 5 min (Fig. 6A), and also summed over a 30-minute observation period (Fig. 6B). Intradermal injection of carvacrol (0.1%, 50 μl) caused significant increases in the accumulated scratching bouts (137.2 ± 33.9) as compared to the control group receiving normal saline (0.9% NaCl, 3.8 ± 1, *n* = 6, *P* < 0.001; Fig. 6B). We then made an intradermal injection of dyclonine into the mouse neck 30 minutes before the injection of carvacrol into the same site. Administration of 50 μl dyclonine at 1, 10 and 50 μM concentrations appreciably reduced the scratching bouts to 130.0 ± 20.3, 82.0 ± 15.0, and 18.0 ± 8.0 (*n* = 6), respectively (Fig. 6B). Hence, dyclonine ameliorates TRPV3 hyperactivity-caused scratching in a concentration-dependent manner.

**Figure 6.**
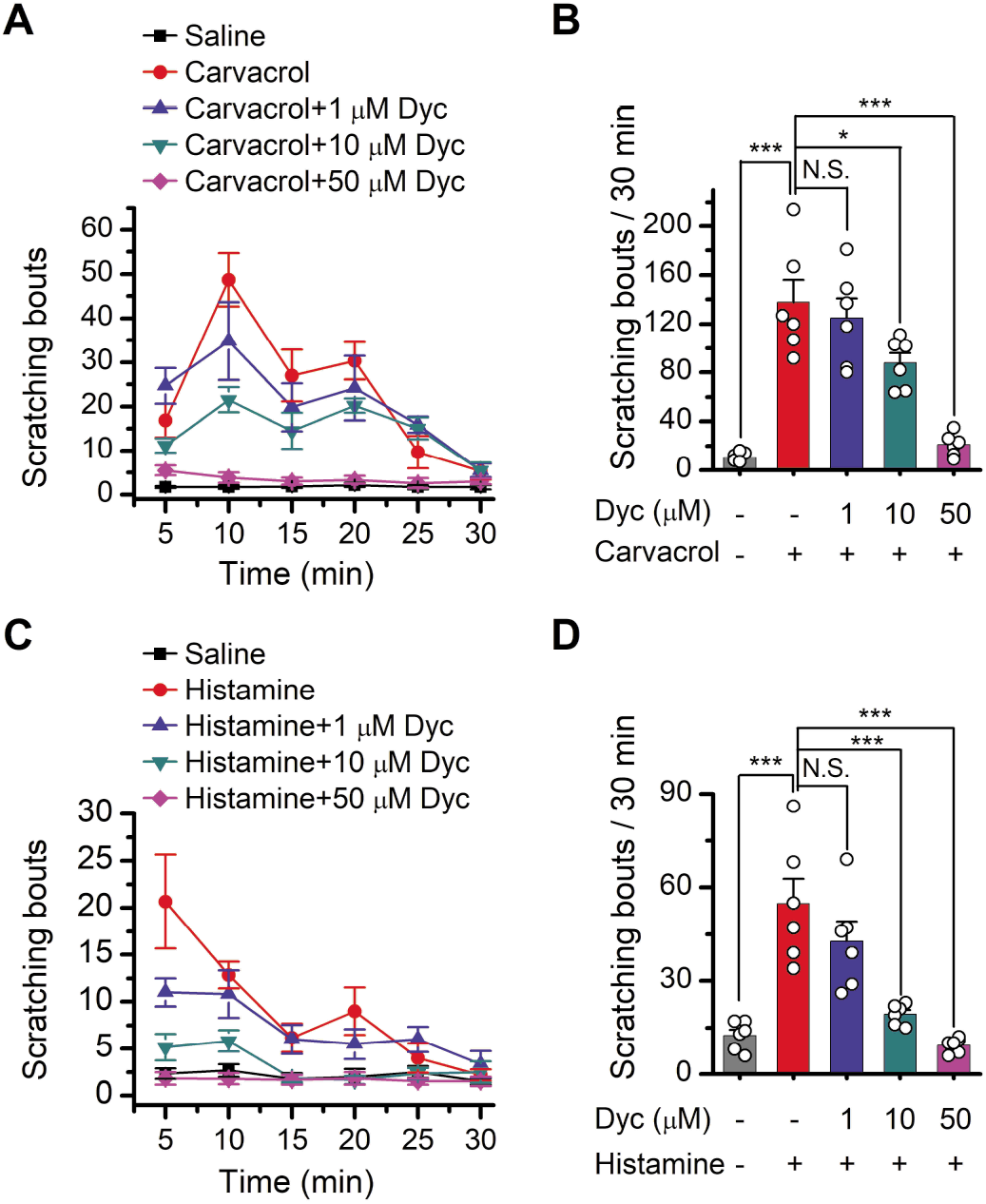
Dyclonine suppresses scratching behavior induced by carvacrol or histamine. **(A)** Summary of the time courses of neck-scratching behaviors in C57BL/6 mice after intradermal injection of 50 μl carvacrol (0.1%) into the mouse neck, with pretreatment with normal saline (0.9% NaCl) containing 0.1% ethanol or different concentrations (1, 10, 50 μM) of dyclonine in the same site. Time for scratching bouts was plotted for each five-minute interval over the 30 minutes observation period. (**B**) Quantification of the cumulative scratching bouts over 30 minutes under different treatments. (*n* = 6; N.S., no significance; \**P* < 0.05; \*\**P* < 0.01; \*\*\**P* < 0.001, by one way ANOVA). (**C**) Time courses of neck-scratching behaviors in response to intradermal injection of 50 μl histamine (100 μM), with pretreatment of normal saline (0.9% NaCl), or different concentrations of dyclonine as indicated. (**D**) Summary plots of the number of scratching bouts over 30 minutes under different treatments as indicated. (*n* = 6; N.S., no significance; \**P* < 0.05; \*\**P* < 0.01; \*\*\**P* < 0.001, by one way ANOVA).

Histamine is another classic inducer of pruritus and has been used to induce experimental itch for decades (Melton and Shelley, 1950). We further examined whether dyclonine could alleviate histamine-evoked acute itch. As illustrated in Fig. 6C-D, intradermal injection of histamine (100 μM) elicited a remarkable increase in the number of scratching bouts (accumulated number = 48.2 ± 10.8 vs. 11.5 ± 1.7 of the saline control, *n* = 6, *P* < 0.001). As above, pretreatment with 50 μl dyclonine (1, 10 and 50 μM) significantly suppressed the number of scratching bouts to 40.6 ± 7.3, 17.5 ± 5.5, and 8.2 ± 3.1 (*n* = 6), respectively. This observation, therefore, corroborates the anti-itch/scratching effect of dyclonine.

### Effects of dyclonine on single TRPV3 channel activity

We then examined the functional and molecular mechanisms underlying the inhibition of TRPV3 by dyclonine. To distinguish whether such inhibition arises from the changes in channel gating or conductance, we measured single-channel activity. Single-channel recordings were performed in an inside-out patch that was derived from HEK 293T cells expressing the mouse TRPV3 (Fig. 7). Currents were evoked by 10 μM 2-APB in the absence and presence of dyclonine (30 μM) after sensitization induced by 300 μM 2-APB at a holding potential of either +60 mV or −60 mV (Fig. 7A). To quantify the changes, we constructed all-point histograms and measured the open probabilities and the unitary current amplitudes by Gaussian fitting. We observed that the single-channel open probability was largely decreased by dyclonine from 0.8 ± 0.02 to 0.08 ± 0.01 at −60 mV and from 0.82 ± 0.02 to 0.12 ± 0.01 at +60 mV (*n* = 6), respectively (Fig. 7B). Statistical analysis, however, revealed that dyclonine had no effect on single TRPV3 channel conductance (163.6 ± 6.4 pS v.s. 179.2 ± 5.5 pS for before and after dyclonine treatment; Fig. 7C).

**Figure 7.**
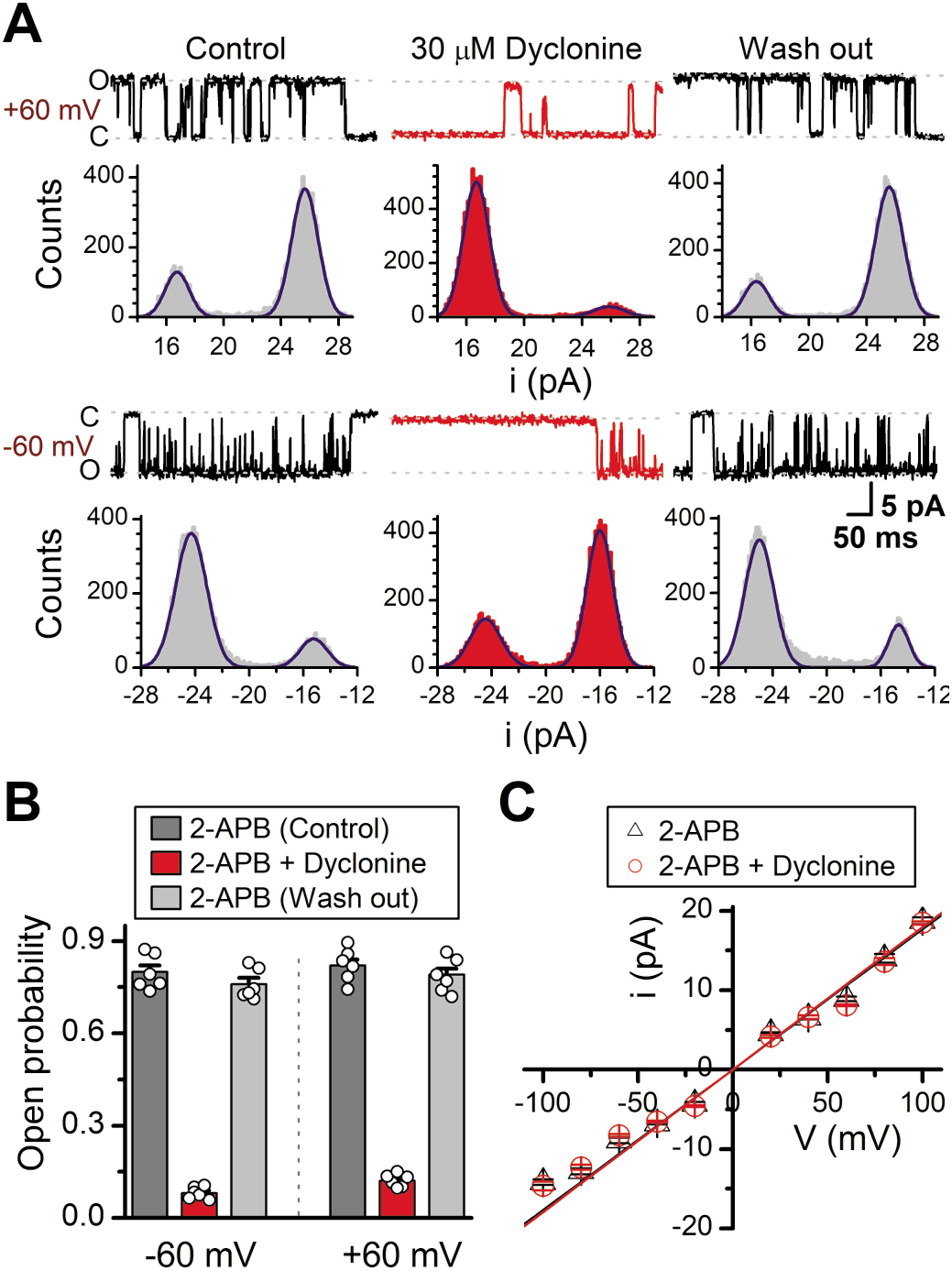
Effects of dyclonine on single channel properties of TRPV3. (**A**) Single-channel currents of TRPV3 were recorded from inside-out membrane patches of HEK 293T cells at two membrane potentials (±60 mV) in symmetrical 150 mM NaCl and were low-pass filtered at 2 kHz. Currents were evoked by 10 μM 2-APB in the absence and presence of dyclonine (30 μM) after sensitization induced by repetitive 300 μM 2-APB. All-point amplitude histograms of single-channel currents were shown below the current traces. The histograms were fit to sums of two Gaussian functions to determine the average amplitudes of currents and the open probabilities. Dotted lines indicate the opened channel state (O) and the closed channel state (C), respectively. (**B**) Summary of effects of dyclonine on TRPV3 single-channel open probability. Dyclonine (30 μM) decreased TRPV3 open probability from 0.8 ± 0.02 to 0.08 ± 0.01 at −60 mV (*n* = 6), and from 0.82 ± 0.02 to 0.12 ± 0.01 at +60 mV (*n* =6), respectively. (**C**) I-V relationships of TRPV3 single-channel current evoked by 10 μM 2-APB without (black triangles) and with 30 μM dyclonine (red circles). Unitary conductance measured by fitting a linear function were 163.6 ± 6.4 pS and 179.2 ± 5.5 pS for before and after treatment by dyclonine, respectively.

### Dyclonine is an open-channel blocker

In order to understand the molecular mechanism underlying the blockade of TRPV3 by dyclonine, we utilized the molecular docking approach to model their interaction. Fig. 8A illustrates the optimized structure of TRPV3-dyclonine complex generated by docking and dynamic simulations. The principal interactions derived from the refined complex structure were analyzed using the Induce-Fit-Docking program (Sherman et al., 2006). The best receptor–ligand complex was evaluated by the extra precision scoring. Three possible docking boxes designated as pocket A, B and C are shown in Fig. 8A. Expanded views of the docking boxes show that three residues L655, I674 and G638 were located in the center of each binding pocket (Fig. 8B). To further delineate dyclonine-interacting residues, we systematically mutated the suspected residues in the cavity of TRPV3 channel to alanine. Among all ten mutants of TRPV3, mutations L655A and F666A greatly reduced the inhibitory effect of dyclonine, whereas the mutants Y622A, L642A and I659A showed higher sensitivity to dyclonine than wild-type channel (Fig. 8C-D). Indeed, substitution of the residue by alanine caused up to ~100 folds change in the inhibition potency of dyclonine. The dose-response curves were fitted with a Hill equation, and the corresponding IC_50_ values for each TRPV3 mutant were as follows: IC_50_ = 282.4 ± 33.4 μM for L655A; IC_50_ = 414.7 ± 23.9 μM for F666A; IC_50_ = 0.39 ± 0.04 μM for Y622A; IC_50_ = 0.25 ± 0.02 μM for L642A and IC_50_ = 0.56 ± 0.06 μM for I659A, compared to IC_50_ = 3.2 ± 0.24 μM for WT TRPV3 (Fig. 8D-E). However, F666 is located below the upper filter and behaves with a bulky hydrophobic side chain, which may play a role in maintaining the shape of this pocket at the open state. This may be the reason why F666A is capable of decreasing the inhibition of dyclonine. Notably, all ten mutant channels were functional and produced robust responses to 2-APB (Fig. 8F). These data suggest that dyclonine interacts with the pore cavity of TRPV3 channel, thereby preventing ion passage and resulting in channel inhibition.

**Figure 8.**
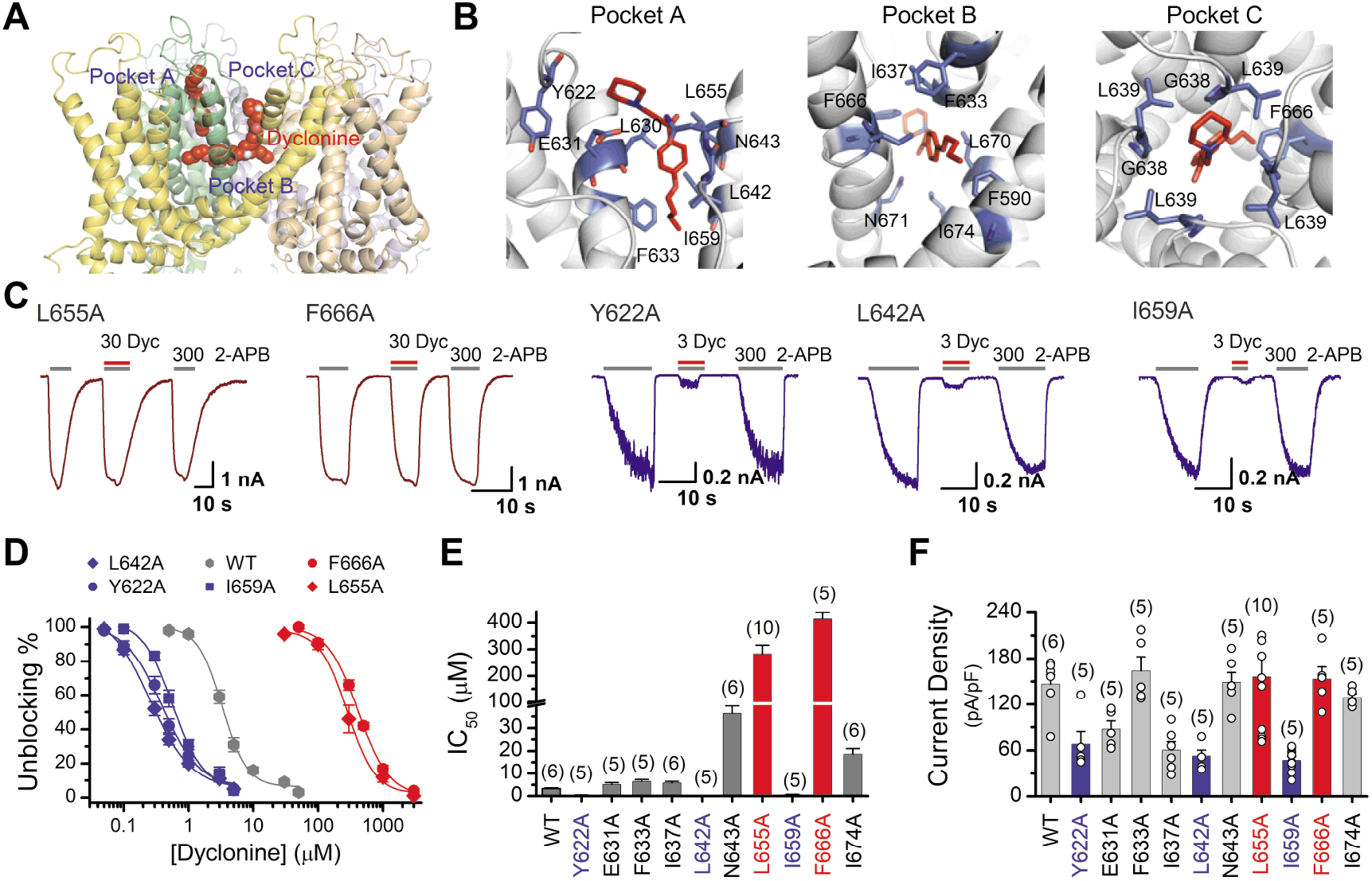
Molecular residues involved in dyclonine-TRPV3 interaction. (**A**) Overall view of the mTRPV3-dyclonine complex. Three putative binding pockets for dyclonine in the pore cavity of mTRPV3 channel (PDB ID code: 6DVZ) are denoted as A, B and C. Four subunits of the tetramer are distinguished by different colors, and dyclonine in a schematic structure is shown in red. (**B**) Detailed binding modes of dyclonine in each binding pocket and the putative interaction residues are labeled. Dyclonine is displayed in sticks for emphasis. (**C**) Representative whole-cell recordings show reversible blocking of 2-APB (300 μM)-evoked responses by dyclonine (30 or 3 μM) in HEK 293T cells expressing mutant TRPV3 channels as indicated, respectively. The combination of 30 or 3 μM dyclonine and 2-APB was applied following the control currents evoked by a saturated concentration of 2-APB (300 μM, initial grey bar). Holding potential was −60 mV. Bars represent duration of stimuli. (**D**) Concentrationresponse curves of dyclonine on inhibition of the TRPV3 mutants. Solid lines represent fits by a Hill equation, with the half-maximal inhibitory concentration (IC_50_) shown in (**E**). For comparison, the dose-response curve of wile-type channel is displayed in gray. Two point mutations (L655A and F666A) reduced the inhibitory efficiency of dyclonine, while the other three point mutations (Y622A, L642A and I659A) enhanced the inhibitory effects of dyclonine on TRPV3 currents. (**F**) Average current responses of mutant channels compared with wild-type TRPV3 channels. Each substitution of putative residues by alanine retained their normal responses to 2-APB. Numbers of cells are indicated in parentheses.

## Discussion

As a multimodal sensory channel, TRPV3 is abundantly expressed in keratinocytes and implicated in inflammatory skin disorders, itch, hair morphogenesis, and pain sensation (Broad et al., 2016). Human Olmsted syndrome has been linked to the gain-of-function mutations of TRPV3 (Lai-Cheong et al., 2012; Lin et al., 2012; Agarwala et al., 2016). Synthetic and natural compounds, like isopentenyl pyrophosphate (Bang et al., 2011), 17(R)-resolvin D1 (Bang et al., 2012), forsythoside B (Zhang et al., 2019), diphenyltetrahydrofuran osthole (Higashikawa et al., 2015) and ruthenium red (Xu et al., 2002) have been proposed to inhibit TRPV3 channels. Due to either or both the lack of targeting specificity and the clinical application, their remedial potential remains to be determined. Hence, identifying and understanding clinical pharmaceutics that selectively target TRPV3 channels will help to conceive therapeutic interventions.

Dyclonine is a topical antipruritic agent and has been used for clinical treatment of itching and pain for decades (Greifenstein et al., 1956; Gargiulo et al., 1992). While the therapeutic effect of dyclonine has been attributed to the inhibition of cell depolarization, the underlying mechanisms have not been fully understood. In the present study, we provide several tiers of evidence that dyclonine selectively inhibits TRPV3 channel. Such inhibition was observed for TRPV3 responses to both chemical and thermal activation, suggesting dyclonine is a condition-across inhibitor. Accordingly, dyclonine efficiently blocked the excessive activation of TRPV3 mutants and prevented cell death. Single-channel recordings revealed that dyclonine effectively suppresses the channel open probability without changing single-channel conductance. These data not only supplement a molecular mechanism for the therapeutic effect of dyclonine, but also suggest its application to curb TRPV3-related disorders. Using mouse model, we indeed observed that dyclonin ameliorates the TRPV3 hyperactivity-caused itch/scratching behaviors, indicating its therapeutic inhibition effect being maintained in vivo. As TRPV3 responds to moderate temperatures (30 - 40 °C), dyclonine may thus be used to alleviate skin inflammations persisted in physiological temperatures. Also, as a clinical drug dyclonine has been widely used and thus has shown its safety to human body. In addition, as a selective and potent inhibitor, dyclonine can also be a research tool to dissect the physio- and pathological characteristics of TRPV3 channel.

The current data also unraveled the molecular residues within TRPV3 channel pore that regulate the effect of dyclonine. Dyclonine interacts likely with L655 and F666 located in the pocket A and C, and thus regulates the ion-conducting pathway. In reality, since binding domains A and C are arranged one by one along the ion-conducting pathway, it is unlikely that dyclonine can occupy these two sites at the same time. Interestingly, the size of pocket A is distinct in the *apo*/resting and open states. Likely, binding of dyclonine into this pocket could prevent the structural rearrangements of pore loop during TRPV3 gating. Since F666 is located below the filter and behaves with a bulky hydrophobic side chain, it may maintain the shape of this pocket at the open state thereby hindering the inhibition effect of dyclonine. Moreover, we identified Y622, L642 and I659 residues whose mutation upregulates the inhibitory effect of dyclonine. These molecular sites could thus be targeted for developing therapeutics for chronic pruritus, dermatitis and skin inflammations.

## Material and Methods

### cDNA constructs and transfection in HEK 293T cells

The wild-type mouse TRPV3 and human TRPV3 cDNAs were generously provided by Dr. Feng Qin (State University of New York at Buffalo, Buffalo, USA). The GFP-mTRPV3 wild-type and the mutants (mTRPV3-G573S and mTRPV3-G573C) in pcDNA4/TO vector were gifts from Dr. Michael X. Zhu (The University of Texas Health Science Center at Houston, Houston, USA). The wild-type frog TRPV3 was kindly provided by Dr. Makoto Tominaga (Department of Physiological Sciences, SOKENDAI, Okazaki, Japan). All mutations were made using the overlap-extension polymerase chain reaction (PCR) method as previously described (Tian et al., 2019). The resulting mutations were then verified by DNA sequencing. HEK 293T and T-Rex 293 cells were grown in Dulbecco’s modified Eagles medium (DMEM, Thermo Fisher scientific, MA, USA) containing 4.5 mg/ml glucose, 10% heat-inactivated fetal bovine serum (FBS), 50 units/ml penicillin, and 50 mg/ml streptomycin, and were incubated at 37°C in a humidified incubator gassed with 5% CO_2_. For T-Rex 293, blasticidin S (10 μg/ml) was also included. Cells grown into ~80% confluence were transfected with the desired DNA constructs using either the standard calcium phosphate precipitation method or lipofectamine 2000 (Invitrogen) following the protocol provided by the manufacturer. Transfected HEK 293T cells were reseeded on 12 mm round glass coverslips coated by poly-L-lysine. Experiments took place ~12-24 h after transfection.

### Mouse epidermal keratinocyte culture

The animal protocol used in this study was approved by the Institutional Animal Care and Use Committee of Wuhan University. Primary mouse keratinocytes were prepared according to the method previously described (Pirrone et al., 2005; Luo et al., 2012). Briefly, newborn wild-type C57B/6 mice (postnatal day 1–3) were deeply anaesthetized and decapitated and then soaked in 10% povidone-iodine, 70% ethanol, and phosphate-buffered saline for 5 min, respectively. The skin on the back was removed and rinsed with pre-cold sterile phosphate-buffered saline (PBS) in a 100-mm Petri dish and transferred into a 2-ml tube filled with pre-cold digestion buffer containing 4 mg/ml dispase II and incubated overnight at 4 °C. After treatment with dispase II for 12-18 h, the epidermis was gently peeled off from dermis and collected. Keratinocytes were dispersed by gentle scraping and flushing with KC growth medium (Invitrogen, Carlsbad, CA). The resulting suspension of single cells was collected by centrifuge and, cells were seeded onto coverslips pre-coated with poly-L-lysine, and maintained in complete keratinocyte serum-free growth medium (Invitrogen, Carlsbad, CA). Cell culture medium was refreshed every two days. Patch-clamp recordings were carried out 48 h after plating.

### Electrophysiological recording

Conventional whole-cell and excised patch-clamp recording methods were used. For the recombinant expressing system, green fluorescent EGFP was used as a surface marker for gene expression. Recording pipettes were pulled from borosilicate glass capillaries (World Precision Instruments), and fire-polished to a resistance between 2–4 MΩ when filled with internal solution containing (in mM): 140 CsCl, 2.0 MgCl_2_, 5 EGTA, 10 HEPES, pH 7.4 (adjusted with CsOH). Bath solution contained (in mM): 140 NaCl, 5 KCl, 3 EGTA, and 10 HEPES, pH 7.4 adjusted with NaOH. For recordings in keratinocytes, the bath saline consisted of (in mM) 140 NaCl, 5 KCl, 2 MgCl_2_, 2 CaCl_2_, 10 glucoses, 10 HEPES, pH 7.4 adjusted with NaOH and the pipette solution contained (in mM): 140 CsCl, 5 EGTA, and 10 HEPES, pH 7.3 adjusted with CsOH. For single-channel recordings, the pipette solution and bath solution were symmetrical and contained (in mM) 140 NaCl, 5 KCl, 3 EGTA, 10 HEPES, pH 7.4. Isolated cells were voltage clamped and held at −60 mV using an EPC10 amplifier with the Patchmaster software (HEKA, Lambrecht, Germany). For a subset of recordings, currents were amplified using an Axopatch 200B amplifier (Molecular Devices, Sunnyvale, CA) and recorded through a BNC-2090/MIO acquisition system (National Instruments, Austin, TX) using QStudio developed by Dr. Feng Qin at State University of New York at Buffalo. Whole-cell recordings were typically sampled at 5 kHz and filtered at 1 kHz, and single-channel recordings were sampled at 25 kHz and filtered at 10 kHz. The compensation of pipette series resistance and capacitance were compensated using the built-in circuitry of the amplifier (>80%) to reduce voltage errors. Exchange of external solution was performed using a gravity-driven local perfusion system. As determined by the conductance tests, the solution around a patch under study was fully controlled by the application of a flow rate of 100 μl/min or greater. Dyclonine hydrochloride, carvacrol and histamine were purchased from MCE (Med Chem Express). Unless otherwise noted, all chemicals were purchased from Sigma (Millipore Sigma, St. Louis, MO). Water-insoluble reagents were dissolved in pure ethanol or DMSO to make a stock solution and diluted into the recording solution at the desired final concentrations before the experiment. The final concentrations of ethanol or DMSO did not exceed 0.3%, which had no effect to currents. In the scratching behavior experiments, carvacrol was firstly dissolved in 10% ethanol and then diluted in normal saline like histamine before administration. All experiments except those for heat activation were performed at room temperature (22-24°C).

### Ultrafast temperature jump achievement

Rapid temperature jumps were generated by the laser irradiation approach as described previously(Yao et al., 2009). In brief, a single emitter infrared laser diode (1470 nm) was used as a heat source. A multimode fiber with a core diameter of 100 μm was used to transmit the launched laser beam. The other end of the fiber exposed the fiber core was placed close to cells as the perfusion pipette is typically positioned. The laser diode driven by a pulsed quasi-CW current powder supply (Stone Laser, Beijing, China). Pulsing of the controller was controlled from computer through the data acquisition card using QStudio software developed by Dr. Feng Qin at State University of New York at Buffalo. A blue laser line (460 nm) was coupled into the fiber to aid alignment. The beam spot on the coverslip was identified by illumination of GFP-expressing cells using the blue laser during experiment.

Constant temperature steps were generated by irradiating the tip of an open pipette and using the current of the electrode as the readout for feedback control. The laser was first powered on for a brief duration to reach the target temperature and was then modulated to maintain a constant pipette current. The sequence of the modulation pulses was stored and subsequently played back to apply temperature jumps to the cell of interest. Temperature was calibrated offline from the pipette current using the dependence of electrolyte conductivity.

### Cell death analysis by flow cytometry

T-Rex 293 cells were grown in DMEM containing 4.5 mg/ml glucose, 10% (vol/vol) FBS, 50 units/ml penicillin, 50 μg/ml streptomycin, and blasticidin S (10 μg/ml) and were incubated at 37°C in a humidified incubator gassed with 5% CO_2_. Transfections were performed in wells of a 24-well plate using lipofectamine 2000 (Invitrogen). The GFP-TRPV3 (wild-type and G573 mutants) cDNAs in pcDNA4/TO vector were individually transfected into T-Rex 293 cells and treated with 20 ng/ml doxycycline 16 h post-transfection to induce the gene expression following the method as previously described (Xiao et al., 2008). Expression of GFP fluorescence detected by an epifluorescence microscope was used as an indicator of gene expression. After treatments with the compounds for 12 h, cells were collected, washed twice with phosphate-buffered saline (PBS), resuspended and then dyed with propidium iodide (PI, Thermo Fisher Scientific) in the dark according to the manufacturer’s instructions. The membrane integrity of the cells was assessed using a BD FACSCelesta flow cytometer equipped with BD Accuri C6 software (BD Biosciences, USA).

### Evaluation of scratching behavior in mice

To assess itch-scratching behaviors, the hair of the rostral part of the C57B/6 mouse right neck was shaved using an electric hair clipper 24 hours before the start of experiments. Scratching behaviors were recorded on video. The number of itch-scratching bouts was counted through video playback analysis. One scratching bout was defined as an episode in which a mouse lifted its right hind limb to the injection site and scratched continuously for any time length until this limb was returned to the floor or mouth (Wilson et al., 2013). All behavioral experiments were conducted in a double-blind manner. To examine acute itch induced by carvacrol or pruritogen histamine, mice were firstly placed in an observation box (length, width, and height: 9 × 9 × 13 cm^3^) for acclimatization for about 30 minutes. Then carvacrol (0.1%) or histamine (100 μM) in a volume of 50 μl was injected intradermally into the right side of the mouse neck. To access the effect of dyclonine on itch scratching, normal saline (0.9% NaCl) or dyclonine (1, 10, and 50 μM) was injected intradermally 30 minutes before intradermal injection of carvacrol or histamine (Sun and Dong, 2016; Cui et al., 2018). Behaviors were recorded on video for 30 minutes following the injection of carvacrol or histamine.

### Molecular docking

The molecular docking approach was used to model the interaction between dyclonine and TRPV3 channel protein (PDB ID code: 6DVZ). The 3D structure of dyclonine was generated by LigPrep (Gadakar et al., 2007). Induce-Fit-Docking (IFD) was employed to dock dyclonine into the potential binding pocket using default parameters(Sherman et al., 2006). During this docking process, the protein was fixed while dyclonine was flexible. The best receptor–ligand complex was evaluated by the extra precision scoring and, at last, L655, I674 and G638 were chosen as the center of the docking box, respectively. All structural figures were made by PyMol (http://www.pymol.org).

### Statistics

Data were analyzed offline with Clampfit (Molecular Devices, Sunnyvale, CA), IGOR (Wavemetrics, Lake Oswego, OR, USA), SigmaPlot (SPSS Science, Chicago, IL, USA) and OriginPro (OriginLab Corporation, MA, USA). For concentration dependence analysis, the modified Hill equation was used: Y = A1 + (A2 - A1) / [1 + 10^(logIC_50_ – X)*n_H_], in which IC_50_ is the half maximal effective concentration, and n_H_ is the Hill coefficient. Unless stated otherwise, the data are expressed as mean ± standard error (SEM), from a population of cells (*n*), with statistical significance assessed by Student’s *t*-test for two-group comparison or one-way analysis of variance (ANOVA) tests for multiple group comparisons. Significant difference is indicated by a *p* value less than 0.05 (\**p* < 0.05, \*\**p* < 0.01).

## Acknowledgements

We are grateful to our colleagues and members of Yao lab for comments and discussions, and we also would like to thank the core facilities of College of Life Sciences at Wuhan University for technical help. This work was supported by grants from the National Natural Science Foundation of China (31830031, 31929003, 31671209, 31871174, and 31601864), Natural Science Foundation of Hubei Province (2017CFA063 and 2018CFA016), and the Fundamental Research Funds for the Central Universities.

## Author Contribution

J.Y. designed and supervised the study. Q.L., J.H., X.W., C.P., Y.G., C.X., and J.Y. carried out the experiments and analyzed data. J.W. and Y.Y. performed the molecular docking experiments. D.L. provided technical support and suggestions. Q.L. and J.Y. wrote the paper with inputs from all other authors. All authors discussed the results and commented on the manuscript.

